# Reactive oxygen generation by minimal copper binding peptide motifs

**DOI:** 10.1101/2025.03.26.645443

**Authors:** Vijeesh Vayyattil, Ruchi Sharma, Maciej B. Gielnik, Nina Lock, Magnus Kjaergaard

## Abstract

Intrinsically disordered proteins such as amyloid-β (Aβ), prion protein (PRP) and α-synuclein bind copper-ions through histidine-rich motifs. Several of these copper-binding motifs catalyze formation of reactive oxygen species (ROS), which is believed to be involved in these proteins’ role in neurodegenerative disorders. The catalytic mechanism rely on binding in an ‘in-between state’, which is energetically available to both Cu(I) and Cu(II) and thus allow redox cycling. Presently, our knowledge of ROS generation is based on a few peptides studied for their pathological role and has thus not explored the minimal requirements for this function. Here, we investigate copper coordination and ROS generation in a series of synthetic histidine-rich peptides. We found that a minimal motif of two histidine residues joined by a few glycine residues is sufficient to generate ROS at two and a half times the rate of the corresponding Aβ complex. Copper-coordination and ROS generation are flexible with regard to the spacing of these two histidine residues suggesting that physiologically relevant Cu-binding motifs are degenerate. Spectroscopic studies indicate that ROS activity correlate to coordination by two imidazole nitrogen atoms in a dynamic complex. This study suggests that copper-coordination and ROS production might be more prevalent in intrinsically disordered proteins than realized.

## Introduction

Copper is one of the most common transition metals in the human body, essential for the function of many enzymes and crucial for proper brain function, including synaptic transmission, neuronal development, and signal modulation^1,2^. In complexes, copper primarily exists in two oxidation states: reduced Cu(I) and oxidized Cu(II). The redox activity makes copper an important catalyst involved in electron transfer reactions. However, the side products of the copper redox cycling, particularly hydroxyl radicals produced via the Harber-Weiss reaction, are among the most reactive and damaging forms of reactive oxygen species (ROS)^3–5^.

Amyloid-β (Aβ) peptides form senile plaques during development of the Alzheimer’s disease, the most common human dementia. Aβ isolated from the brain tissue of Alzheimer’s disease patients are found in complex with copper, and purified Aβ binds copper *in vitro*^5–7^. Post-mortem analyses of Alzheimer’s disease patients consistently show oxidative damage in the brain, linking the disease to an imbalance in ROS production or scavenging. Other neurodegenerative diseases that show with signs of oxidative damage are also associated with copper binding polypeptides including α-synuclein in Parkinson’s disease^8^ and prion protein (PrP) in Creutzfeldt-Jakob’s disease^9^.

Efficient ROS production requires that the copper-binding site forms a chemical environment suitable for both oxidation states of copper. This is challenging because Cu(I) coordinates “soft” ligands like sulfur from cysteine or methionine sidechains and prefers linear, trigonal or tetrahedral binding geometry. In contrast, Cu(II) prefers “hard” ligands, such as nitrogen from N-terminal amines, backbone amides, or histidine sidechains, in square planar, pyramidal or bipyramidal geometries^10^. Some enzymes, including blue copper proteins, overcome this problem by utilizing hybrid active sites, which stabilize both redox states, in a single pocket, but each with different geometries^10^.

In contrast to copper-binding enzymes, the polypeptides involved in neurodegenerative disorders are intrinsically disordered. Intrinsically disordered proteins do not form rigid 3D scaffolds into which the metal fits. Instead, metal coordination occurs through short linear motifs, where residues close in the sequence form a metal binding site. Often the peptide becomes locally structured around the metal binding site but does not necessarily form a unique binding geometry. Disordered proteins catalyze redox reactions via a flexible coordination state energetically available for Cu(I) and Cu(II) binding, which is referred to as the “in-between state”. This state acts as a transient intermediate where the redox reaction occurs leading to ROS degeneration^10^. The copper binding polypeptides involved in neurodegenerative disorders such as Aβ, α-synuclein and PrP all have disordered regions that bind metal ions and most likely they all form an “in-between state” leading to ROS formation.

For Aβ peptides, Cu(II) is mainly coordinated in a square-planar geometry by N-terminal amine, a proceeding main-chain carbonyl, and nitrogen atoms of two histidine imidazole sidechains (His6 and His13/14)^11^. On the other hand, Cu(I) binds in a linear geometry involving two of the three histidine residues (His6/13/14) in a dynamic exchange^12^. Variants of Aβ complexed with Cu(II), including Aβ(1-16), Aβ(1-28), Aβ(1-40), and Aβ(1-42), can catalyze ROS production in the presence of ascorbate, with Aβ(1-42) being the most efficient ROS generator^13^. Interestingly, N-terminal truncation of Aβ peptides yields variants such as Aβ(4-x) and Aβ(11-x) with three orders of magnitude higher Cu(II) affinity^14,15^, where x can be either 40 or 42. These truncated forms have amino-terminal copper and nickel (ATCUN) binding motifs, which has the consensus sequence NH^2^-Xxx-Zzz-His, where Xxx and Zzz are any amino acids different than proline^16,17^. ATCUN motifs bind Cu(II) by N-terminal amine, two amides and His imidazole side chain and have been found in many proteins. While ATCUN motifs efficiently complex with Cu(II), they are unable to effectively reduce Cu(II) as they do not bind well to Cu(I) and thus less readily act as redox intermediates^10^. Nevertheless, Aβ(11-x) produce ROS under high ascorbate concentrations^18^ suggesting a more complex coordination than the consensus ATCUN motif.

α-synuclein has two main Cu(II) binding regions: A high affinity N-terminal motif coordinating copper by Met1 amine and amide nitrogen in conjunction with the carboxylate of Asp2^19^; and lower affinity coordinating copper by imidazole nitrogen of His sidechain, the amide nitrogens of Val49 and His50 and a water molecule^20^. Cu(II) can also bind by intermolecular interactions, what can promote oligomerization^21^, Cu(II) might accelerate formation of toxic α-synuclein oligomers and fibrils, which can be rescued by chelating agents^22^. α-synuclein also binds Cu(I) by Met1-Met5 and Met116-Met127 by sulfur atoms of the sulfide groups. Additionally Asp2 carboxylate is also coordinated near N-terminal Cu(I) ion, which can mediate formation of the “in-between state”^23^.

The prion protein (PrP) binds up to six Cu(II) atoms by N-terminal intrinsically disordered region (IDR)^3^. First, up to four copper ions are coordinated by the four octapeptide repeats (residues 60-91 in human PrP) in a concentration-dependent binding mode with high affinity^24–26^. Up to 1:1 Cu(II):octarepeat molar ratio, this region binds a single Cu(II) ion by four imidazole sidechains of His residues, from 1:1 to 2:1 Cu(II):octarepeat molar ratio, each Cu(II) ion is coordinated by two His sidechains and two amide nitrogens^27^, and possibly an additional carbonyl oxygen^24,25^, where starting from above 2:1 Cu(II):octarepeat molar ratio each His sidechain starts to coordinate a single Cu(II) ion with two amide nitrogen and one carbonyl, from the neighboring Gly, binds a single Cu(II) ion^24,25^. Two additional copper ions are coordinated by the amyloidogenic region, compromising residues 90-126, although with a lower affinity^25,26^. Each Cu(II) ion binds in this region independently and each is coordinated by His sidechain and three amide main chain nitrogen atoms or by His sidechain, two amide main chain nitrogen atoms and carbonyl oxygen^26,28^. Interestingly Cu(II) binding to the octarepeat and amyloid regions can induce formation of β-sheet secondary structure, characteristic for disease-related fibrillar structures^29,30^. Like for previous proteins, incubation of PrP with Cu(II) in the presence of ascorbate catalyzes redox cycling^31^. The redox cycling by the Cu(II)-octarepeat domain depends on the copper binding mode, with the cycling quenched at the 1:1 binding mode and active in 4:1 mode, with the amyloidogenic region also producing ROS^32^.

Unlike metalloenzymes, most metal binding IDPs such as Aβ have not evolved under selection to bind copper. The ROS production is thus likely accidental and there is no evolutionary reason to think that copper binding motif characterized in conjunction with neurogenerative disorders are optimized for ROS production. We set out to investigate copper binding and ROS production using minimal copper coordinating peptides. We find that copper complexes with short peptides combining glycine and histidine residues can lead to ROS production that is two and a half fold higher than Aβ_16_. For a series of systematically varied minimal peptides we measure ROS production, binding thermodynamics, reduction potentials and complex geometry. We find that ROS production correlates to a binding mode where two imidazole nitrogen atoms coordinate to the copper-ion leaving the remaining binding sites free for interaction with e.g. oxygen.

## Materials and Methods

*Synthetic peptides:* Synthetic peptides were purchases from Genscript at a purity >95% with the following sequences of (Aβ(1-16) (called Aβ_16_ from here), GHG, GHGHG, GHGHGHG, GHGHGHGHG, GHHG, GHG_2_HG, GHG_3_HG, GHG_4_HG, GHG_5_HG, GHG_6_HG, GHG_7_HG, GFG_2_HG, GYG_2_HG, GWG_2_HG, GMG_2_HG, GDG_2_HG, GEG_2_HG, GQG_2_HG, GNG_2_HG, GKG_2_HG). All peptides except Aβ_16_ were N-terminally acetylated and C-terminally amidated.

### UVVis spectroscopy

The absorption spectra of copper titration into the peptides were studied using a Labbot instrument with a 1 cm quartz cell. Peptide samples at a concentration of 300 μM were prepared in a 50 mM HEPES buffer, pH 6.7. 3 mM CuCl_2_ solutions prepared in the same buffer were titrated into the peptide solution at molar ratios of 0, 1, 2, 3, 4, and 5 using the titration syringe. The absorption spectra were recorded over the spectral range of 450 to 750 nm for each molar ratio after 30 seconds of stirring.

### Isothermal titration calorimetry

All ITC experiments were carried out at a constant temperature of 298 K using a PEAQ-ITC instrument from Malvern Analytical. The metal ions and peptides were dissolved in a 50 mM HEPES buffer with a pH of 6.7. During the experiment, 2 μL injections of a 3 mM CuCl_2 (aq)_ solution (19 injections, 0.4 μL for the first injection only, injection duration: 4 s, injection interval: 150 s) were added to the reaction cell, which contained 200 µM peptide initially. The fitted offset option was used for correcting the heat of dilution during the data analysis. Each titration was repeated three times. A 1:1 binding model was fitted to the thermogram using the Microcal PEAQ-ITC analysis software.

*H*_*2*_*O*_*2*_ *Detection Assay*: The assay was performed using the Cell Technology Fluoro H_2_O_2_ kit. In short: Horseradish peroxidase was dissolved following 20 minutes of sonication and diluted to 10 U/mL and kept at −20 °C. The fluorogenic detection reagent was dissolved in dimethylsulfoxide and stored at −70 °C. A standard curve was generated at 0, 0.25, 0.5, 1.0, 2.0, 4.0, and 8.0 μM H_2_O_2_ and used to convert fluorescence intensity into concentrations. 100 μM peptide and 100 μM CuCl_2_ were incubated at room temperature. Samples were withdrawn after 0, 2, 4, 24, 48 and 72 hours and incubated with enzyme and detection reagent for 10 minutes before detection using a SpectraMax i3 plate reader at 550 nm excitation and 595 nm emission. Background fluorescence of a blank sample was subtracted from each measurement.

### Cyclic voltammetry

Cyclic voltammetry (CV) experiments were performed using an CHI660E potentiostat. The CV measurements were carried out at a scan rate of 100 mV s^-1^, with three scans recorded per peptide-Cu(II) complex solution to study reproducibility/complex stability. An H-cell was used for electrochemical measurements, comprising a glassy carbon electrode (GCE, 1 cm^2^) as the working electrode, a platinum mesh as the counter electrode, and a KCl-saturated Ag/AgCl reference electrode (ElectroCell LF-1). The two chambers in the H-cell was separated with a membrane (Sustainion® 37-50 membrane, dioxide materials). Experiments were conducted in an argon-purged atmosphere to prevent interference from dissolved oxygen. The GCE was polished with 0.05 μm alumina slurry to achieve a mirror finish, followed by 15 minutes of ultrasonication to remove any residue. Electrochemical measurements were performed in 0.1 M potassium phosphate buffer at pH 7.01, with pH adjustments made using concentrated KOH_(aq)_, or HNO3_(aq)_. The peptide concentration was maintained at 0.2 mM, and a ligand-to-Cu(II) ratio of 1:0.9 was used to limit interference from free Cu(II) ions. The Ag/AgCl potential (*V*_Ag/AgCl_) was converted to the potential versus the reversible hydrogen electrode (*V*_RHE_):

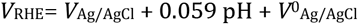

where, *V*^0^_Ag/AgCl_ =0.198 V, *V*_RHE_ is the potential with reference to RHE, and *V*_Ag/AgCl_ is the experimentally measured potential against the Ag/AgCl reference and pH of the phosphate buffer (7.01).

## Results

To study the sequence-function relationships of internal copper-peptide complexes, we studied a series of minimal peptides with between one and four histidine residues. To avoid contributions from other functional groups, we chose glycine as a passive spacer in any other position – allowing only generic interactions with the peptide backbone. Similarly, we used peptides with acetylated N-terminus and amidated C-terminus to avoid interactions with the terminus as occurs in ATCUN motifs. Initially, we focused on peptides with a spacing between histidine residues of a single glycine G(HG)_x_ (x=1-4), as this could in principle allow tetra-dentate binding with two histidine imidazole groups and two backbone nitrogen atoms (Fig. 1). Alternatively, this system could also bind tridentately with a single imidazole and two backbone nitrogen atoms or bivalently with two coordinated imidazole nitrogen atoms. For the longer peptides even more binding modes are available and many different coordination geometries are likely to coexist.

**Fig. 1:**
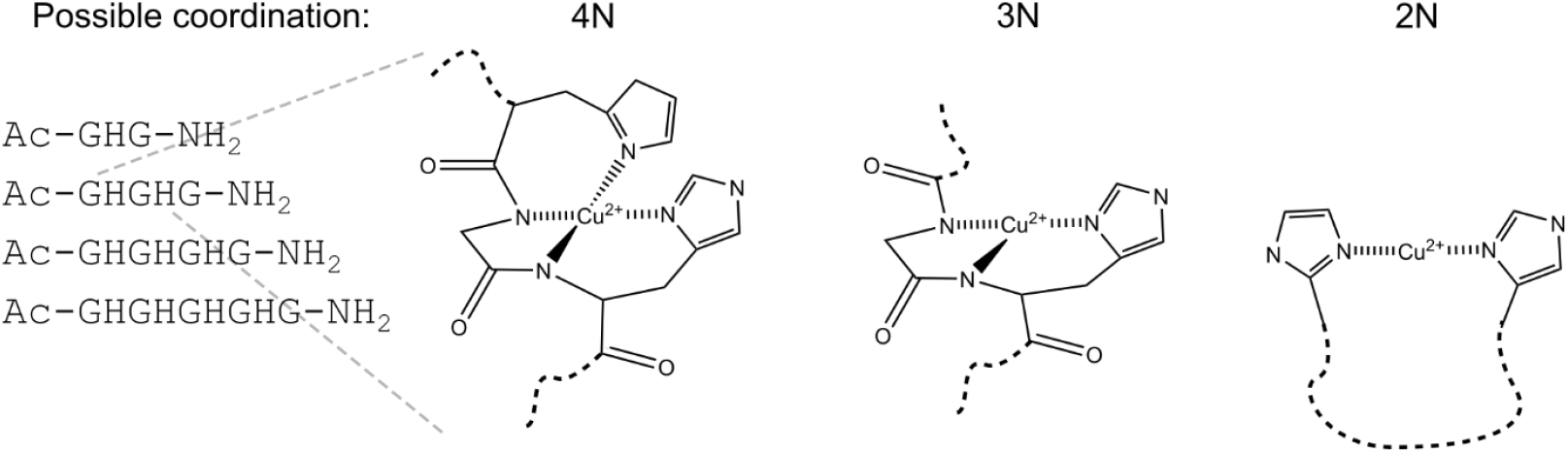
Possible coordination modes of a minimal G(HG)_x_ peptide series. We investigate a minimal peptide series consisting of between one and four histidine residues spaced by single glycine residues. The basic HGH motif could theoretically interact with copper(II)-ions using both imidazoles and two backbone nitrogen atoms (4N), or just via the a single imidazole and two backbone nitrogen atoms. Peptides with two or more histidine residues could also interact with copper(II)-ions in a bidentate manner with a longer flexible linker.

We used a biochemical assay based on peroxidase, where the hydrogen peroxide generated by the copper-complexes are used by peroxidase to oxidize resazurin to resorufin, which can be detected by its fluorescence (Fig. 2A). The catalytic properties of Aβ is recapitulated by Aβ_16_, which contains the metal binding site but do not form fibrils. Neither Aβ_16_ nor copper(II) alone was sufficient to produce a detectable fluorescence, but in complex they steadily formed hydrogen peroxide resulting in accumulation of fluorescence over a period of 72 hours in line with previous reports (Fig. 2B)^13^. For the G(HG)_x_ peptides, the hydrogen peroxide production increased with an increasing number of histidine residues, where peroxide production by the GHGHGHGHG peptide was more than half of what was observed for Aβ_16_ (Fig. 2C). This demonstrates that that ROS production occurs readily in simple copper coordinating peptides.

**Fig. 2:**
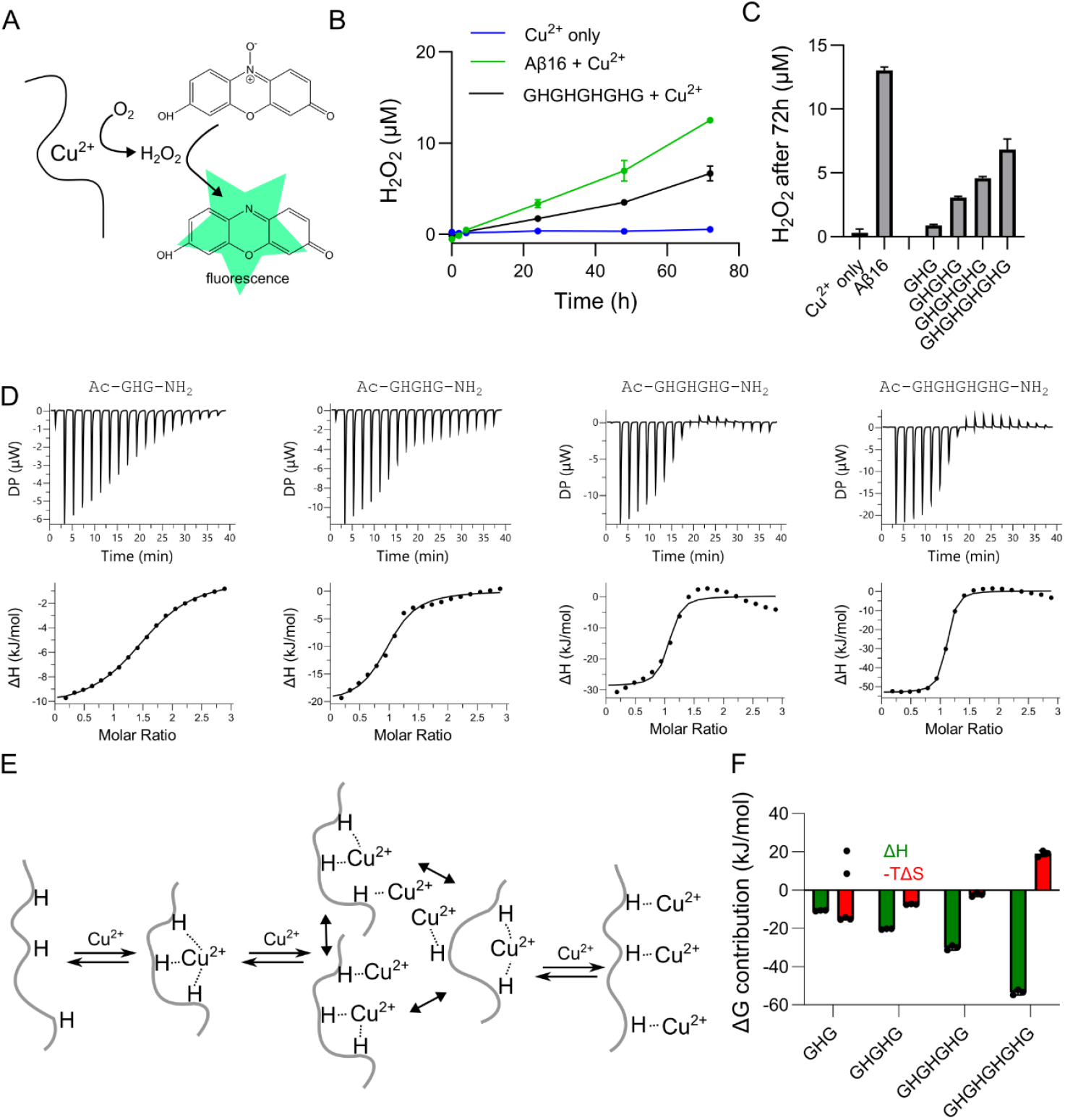
Reactive oxygen species generation by minimal histidine peptides. A) Schematic of fluorescence assay to detect reactive oxygen species. B) Time-courses of the hydrogen peroxide concentrations as determined from fluorescence intensity using a standard curve. C) End-point measurement of hydrogen peroxide for different peptide series relative to the Aβ_16_. D) Isothermal titration calorimetry (ITC) experiments of peptides with different number of histidine residues as a function of the peptide:Cu ratio. The fitted curve represents a 1:1 interaction. Deviations from this model likely stem from complexes with several copper ions bound to the same peptide. E) Decomposition of the enthalpic and entropic components of binding show evidence of enthalpy-entropy compensation.

We used isothermal titration calorimetry to probe the thermodynamics of the interaction of the G(HG)_x_ peptides with copper(II) (Fig. 2D). ITC measurements are complicated by the solubility of copper at the high concentrations required, where copper(II) tends to precipitate unless it is solubilized by weakly chelating buffer components. The binding reaction measured is thus a competitive binding, where the peptide displaces a buffer molecule for coordination to the copper ion. The measured binding constant is thus lower than the actual binding constant and is often referred to as *apparent K*_*D*_ or K_ITC_.^33^ K_ITC_ can be converted to exact binding constant if the binding constant to the buffer is known. However, here the focus was mainly the comparison of motifs, so we compared ITC values recorded under identical conditions (50 mM HEPES, pH 6.7).

The peptide containing a single histidine (GHG) bound Cu^2+^ with an apparent K_ITC_ of 31 µM was well-described by a single-site model (Fig. 2D, Fig. S1, Table S1). The peptide with two histidine residues (GHGHG) showed a reproducible deviation from a one-site model at higher concentrations. The data was better described by a fit to a two-site model with K_ITC_ values in the high nanomolar and mid-micromolar ranges (Fig. S2, Table S2). We ascribed it to initial formation of a 1:1 complex that gradually got displaced by a 1:2 complex (peptide:Cu). Due to large number of free fitting parameters in such models, there was considerable uncertainty in the fitted parameters.

The binding curve for peptides with three or four histidine residues were noticeably biphasic with an initial exothermic reaction followed by an endothermic reaction (Fig. 2D). This suggested that the initially bound state was replaced by a complex with a lower enthalpy of binding resulting in a net endothermic reaction. The saturation was noticeably steeper indicating a higher affinity, which was also reflected in the K_ITC_ for the GHGHGHGHG peptide of around 1 µM. The data is better described by two-, three-site sequential models, however again with substantial uncertainties about the fitted parameters (Fig. S3-4, Table S3,4).

The complexity of the binding reaction was expected as several binding modes can be envisioned for as illustrated in Fig. 2E for the GHGHGHG peptide. At low stoichiometries, 1:1 binding dominated, whereas higher stoichiometries will resulted in increasing amounts of 1:2 and eventually 1:3 binding (peptide:Cu). This multi-state binding model explains the endothermic phase as the states coordinating one or two histidine residues have a lower enthalpy of binding than those with three or four histidine residues coordinated. It is unrealistic to uniquely define the binding mode from these data due to covariance between fitting parameters. A single site model fitted the first part of the binding reaction of most peptides reasonably well (the GHGHGHG peptide being the exception). While neither of these fits are perfect, they at least allow estimation of the thermodynamic parameters of the binding reaction.

The enthalpy of binding increased steadily with the number of histidine residues from ~10 to ~60 kJ/mol. This parameter could be estimated directly from the magnitude of peaks in the thermogram (Fig. 2D) and were thus the most robustly determined parameters regardless of the binding model. The increase in enthalpy was partially offset by a negative entropic contribution (Fig. 2F) – likely from the loss of conformational freedom in the peptide when multiple histidine residues are tied to the same metal-ion. This suggested that the affinity enhancement from multi-dentate binding to metal-ions is limited by enthalpy-entropy compensation.

### Spacing of coordinating residues

Next, we wanted to investigate the role of spacing between coordinating histidine residues. To minimize the complication of binding stoichiometries above 1:1, we focused on peptides with just two histidine residues separated by between zero to seven glycine residues (Ac-GHG_x_HG-NH_2_, x = 0-7) (Fig. 3A). The GHG_x_HG peptides were screened for peroxide production as described above (Fig. 3B). The GHHG peptide contained two adjacent histidine residues that form the core of the copper binding motif in Aβ. GHHG and Aβ_16_ had similar catalytic efficiency suggesting that two adjacent histidine residues were sufficient to form the minimal catalytic motif of Aβ. As reported in Fig. 2, the GHGHG peptide had a markedly lower catalytic efficiency than Aβ_16_ suggesting that inclusion of a single-glycine spacer induced a binding geometry that was less favorable for catalysis. The highest catalytic efficiency was seen for the GHGGHG peptide, which had a catalytic activity two and a half times that of Aβ_16_. Longer spacings appeared similar with approximately twice the catalytic efficiency of Aβ. This suggests that ROS production was favored by a coordination mode favored by spacings two or greater but did not require a specific spacing suggesting that the peptide chain retained some flexibility in the complex.

**Fig. 3:**
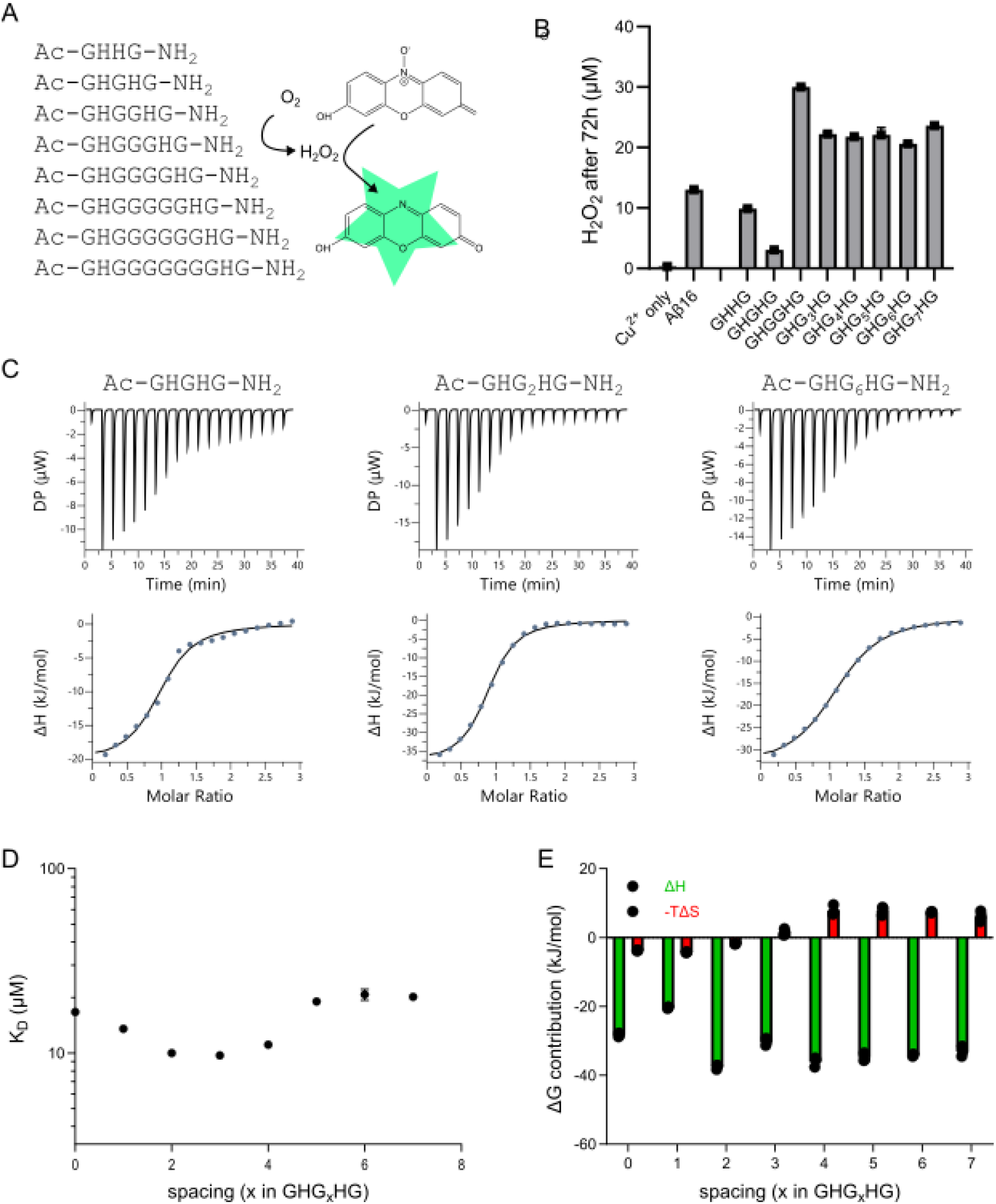
Simple Cu^2+^-coordination motifs exhibit a higher ROS production than Aβ_16_. A) Series of synthetic peptides varying the spacing between histidine residues. The histidine coordination is likely heterogeneous, and the proposed model is one plausible model for the peptides with longer spacers. B) ROS production from peptide:copper complexes evaluated using a fluorescent reporter assays show the most active peptide has 3-fold higher activity than Aβ_16_. C) ITC experiments for peptides binding to copper(II). The fit represents a 1:1 model. D) Spacing dependence of the apparent K_D_ from ITC data. The data are not corrected for competition with buffer components and are thus only apparent values for internal comparison. E) Enthalpic and entropic contributions to the binding energy from the one-site fit suggest that short spacings lead to an enthalpic penalty and longer spacings an entropic penalty.

We performed binding experiments by ITC for each of the peptides in the GHG_x_HG series (Fig. 3C). Each peptide showed saturable, exothermic binding to Cu^2+^, can be attributed to 1:1 binding as above. There were noticeable deviations from the 1:1 model at short spacings as discussed above, suggesting displacement of 1:2 binding modes at higher stoichiometries. With increasing spacing, the binding curve is increasingly well-fitted by a 1:1 binding model (Fig. 3C). The affinity shows a non-monotonic dependence on the number of residues separating the two coordinating residues with a minimal K_D_ for x = 2-3 (Fig. 3D). For peptides with spacing of more than 3 residues, the affinity drops with spacer length. The series could not be extended further than x=7 as it is increasingly more difficult to synthesize and purify highly repetitive peptides. However, while there was a trend in the affinity as function of spacing, the amplitude of this change was modest with little more than a factor of two difference between the strongest and weakest K_D_.

To understand how histidine spacing affects catalysis, we decomposed affinities into enthalpic and entropic components (Fig. 3E). The entropic component starts out favorable but gets increasingly unfavorable with increasing spacing. This is expected as the entropic cost of forming a complex increased with increasing spacing. The enthalpic contribution has a non-monotonous dependence on spacing with a minimum at x=1 (Fig. 3E), which can also be seen directly in the raw thermogram (Fig. 3C). The enthalpy is highly variable at spacing of three or less residues suggesting that the conformational strain of closely spaced residues affect the geometry of the binding site and thus the energetics. At spacings of four or more residues the energetics of binding are relatively stable. Overall, our results suggest that the relatively low dependence of the binding affinity on histidine spacing is in part due to enthalpy-entropy compensation. Intriguingly, the catalytic activity seems to correlate to a favorable enthalpy of binding. This again suggests that the peroxide formation is sensitive to the coordination of the Cu-ion by the peptide.

### Alternative coordinating residues

Next, we investigated whether two coordinating histidine residues were necessary for formation of peroxide or whether other residues could replace histidine. Using the GHGGHG peptide as baseline, we exchanged the first histidine residue (GXGGHG) with plausible copper coordinating side chains and a few non-plausible residues as control (Fig. 4A). For each of these residues, we tested the ROS generation. The only amino acid in this series that led to measurable ROS production was cysteine (Fig. 4B). For each peptide, we performed also ITC and fitted a 1:1 binding model to the thermogram (Fig. 4C). All other peptides except for GCGGHG were well fitted by a 1:1 model. The GCGGHG peptide displayed a strongly exothermic reaction that did not resemble the profile of a binding reaction (Fig. S5) – possibly due to redox reactions involving the thiol group in the cysteine. The best binding was observed for the histidine containing peptide as expected, whereas slightly weaker affinity was observed for amide (Q, N) or amine (K) containing side chain, whereas carboxy (D,E), thioether (M) or aromatic (F,Y,W) functional groups hardly enhanced affinity compared to the single histidine containing peptide (Fig. 4D). For all other peptides, the thermodynamics of the binding resembled the GHG peptide, suggesting that the single histidine is the dominant binding interaction (Fig. 4E). In conclusion, this shows that only the strongly coordinating side chains from histidine and cysteine lead to ROS production, and for efficient ROS production copper has to be coordinated by at least two side chains.

**Fig. 4:**
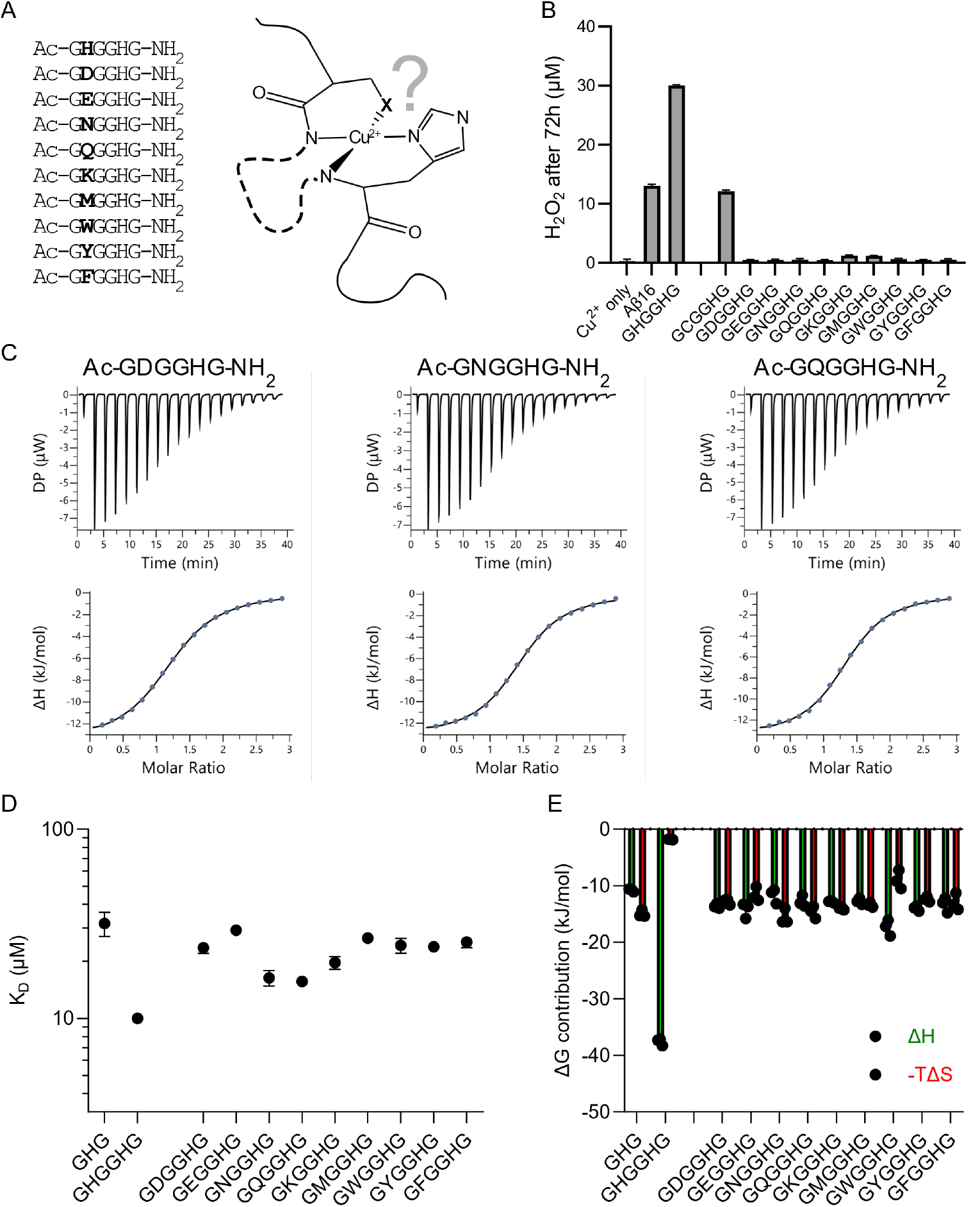
Alternative coordinating residues in Ac-GXGGHG-NH_2_ peptides do not lead to ROS production. Selected isothermal titration calorimetry thermograms of peptides with two histidine residues separated by a various number of glycine residues. The experiments are done under identical conditions (200 µM peptide, 3 mM CuCl_2_). The fitted curve represents a one-site model. B) Dissociation constant from one-site fit to ITC data. Error bars represent s.d. from three repeats and are in some cases hidden under the symbol. C) Decomposition of enthalpic and entropic contributions to the free energy of binding from ITC experiments.

### Redox potential assessed by cyclic voltammetry

ROS production involves cycling between Cu(I) and Cu(II) states, so we reasoned that the difference in ROS production between the different Cu:peptide complexes might be related to the energetics of this transition. We employed cyclic voltammetry to investigate this redox process in the GHG_x_HG peptide series, where the Cu(II) undergoes an initial electrochemical reduction to Cu(I), followed by a subsequent oxidation back to Cu(II) (Fig. 5A). Initial experiments were conducted with a wider potential window resulting in formation of both metallic copper and Cu(III). However, the resulting voltammograms also exhibited signatures of irreversible redox processes. To ensure the selective observation of the reversible Cu(I)/Cu(II) redox couple, the potential window was subsequently narrowed. Nevertheless, some peptide complexes (e.g. GHG_7_HG) still exhibit electrochemical signatures indicative of Cu(III) species (Fig. 5B), but this peptide is an outlier in the series (Fig. 5C), likely due to a less defined redox transition. The oxidation profile of the GHGHG peptide has a clear additional peak at a potential of ~0.5 V, which corresponds to the oxidation potential of free Cu^+^ (Fig. 5A). A shoulder is seen at this potential in the voltammogram of several peptide complexes suggesting the presence of a small amount of free Cu^+^. Despite these complications, the cyclic voltammetric experiments allow determination of oxidation and reduction potentials of the Cu(I)/Cu(II) redox couple in all peptide:copper complexes (Fig. 5C).

**Fig. 5:**
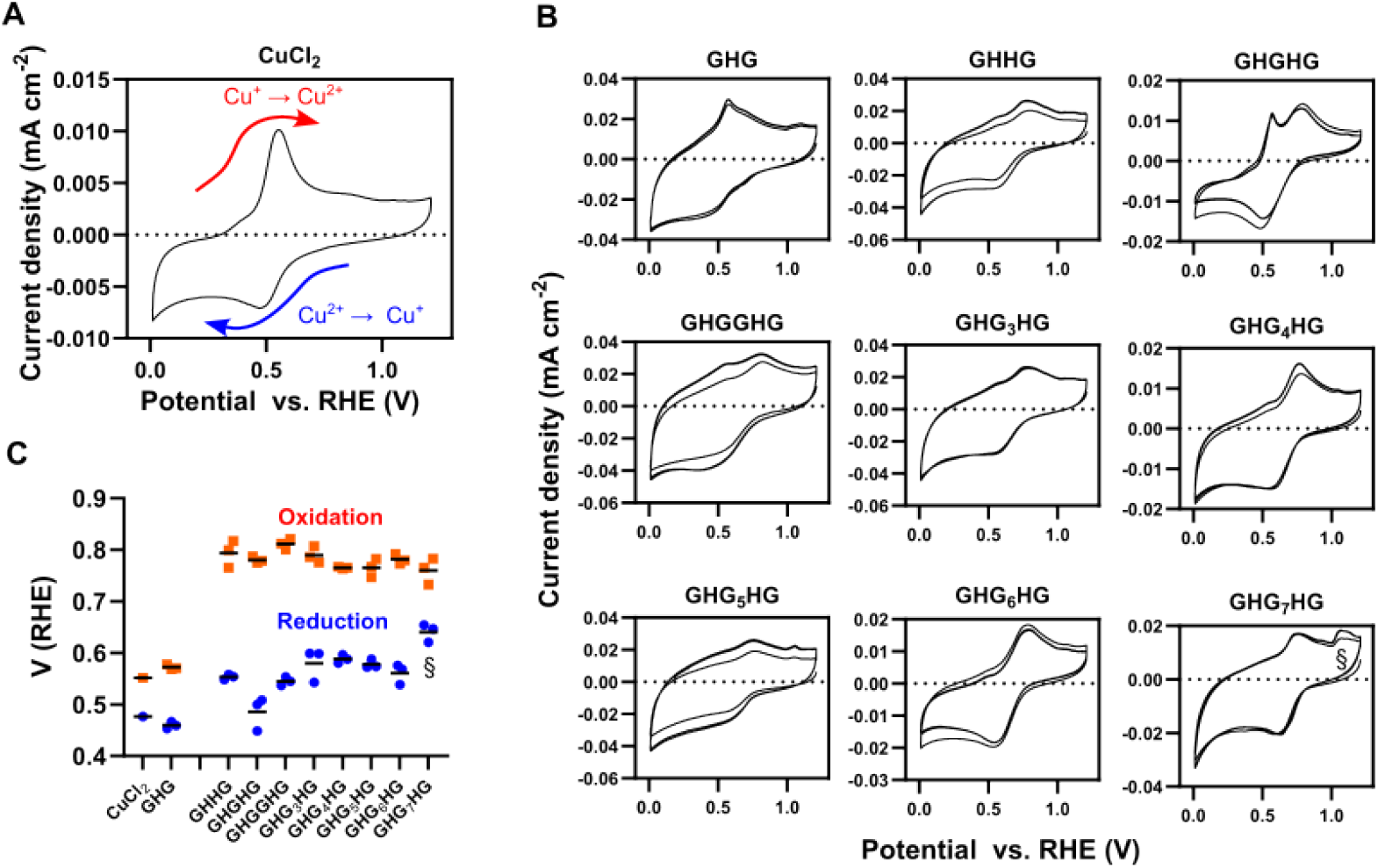
Reduction and oxidation potentials for Cu:peptide complexes from cyclic voltammetry (CV). A) Annotated CV curve from CuCl_2_ with clear peaks for Cu^2+^ to Cu^+^ reduction and Cu^+^ to Cu^2+^ oxidation. B) CV curves of peptide copper complexes (0.2 mM) all show shifts in maxima suggesting that the peptide bind and stabilize the copper ion. The GHG peptide shows a peak at the position of 0.57 V indicative of free copper suggesting that not all copper is bound. C) Oxidation and reduction potentials extracted from the maxima and minima of the current density in the CVs from three independently prepared samples. Differences in amplitude could be from the slight variation in concentration of peptide copper complex at electrode surface at each repeat or a slight modification of the complex structure after initial reduction and oxidation. § suggests signs of formation of Cu^3+^, which subsequently complicate accurate determination of the reduction trace.

**Fig. 6:**
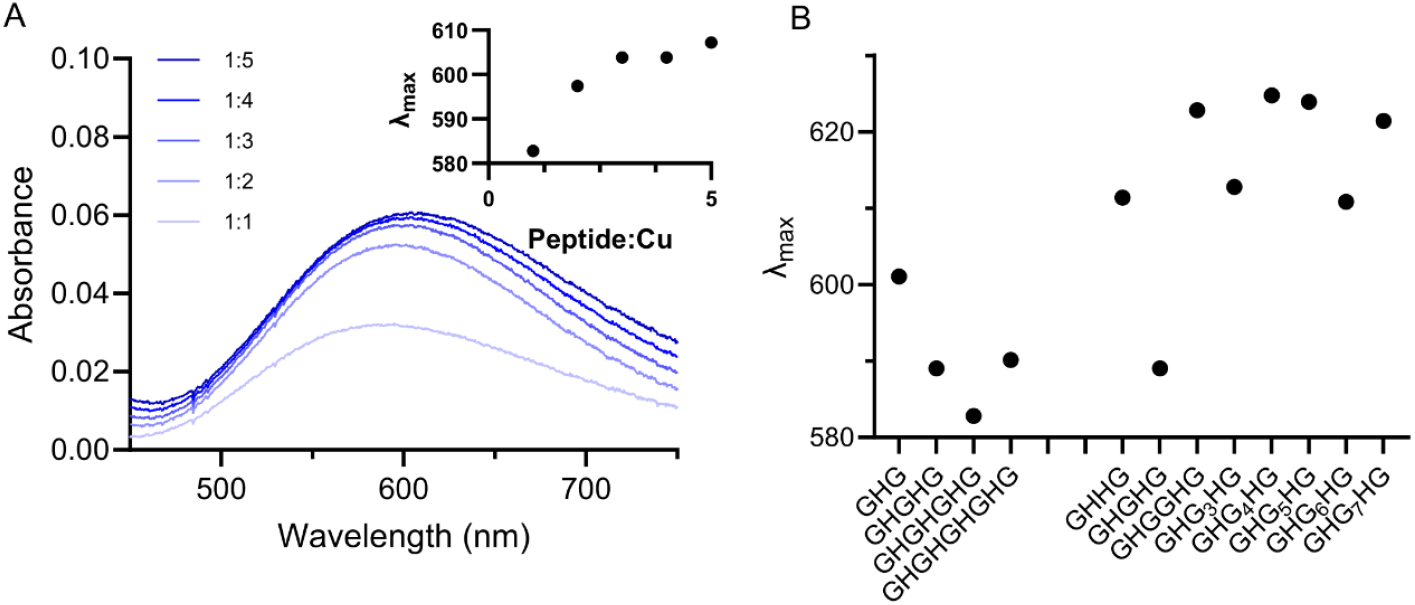
UV-Vis spectra of peptide copper complexes. A) Titration of GHGHGHG peptide with increasing amounts of copper. (insert) Emission maximum as a function of peptide:copper ratio. B) Absorption maxima of 1:1 complexes of peptide:copper complexes.

The oxidation potentials increase ~0.25 V between peptides with one and two histidine residues suggesting an interaction and stabilization of this species. In contrast, there was little variation in oxidation potential dependence in the GHG_x_HG-series suggesting that the exact spacing is not crucial for the Cu^+^ complexes. In the reduction profile, the GHGHG peptide only marginally increased the reduction potential whereas the remaining peptides increased the reduction potential between 0.1-0.15 V relative to CuCl_2_. This mirrors the pattern observed by ITC, where the GHGHG leads to a much weaker binding than the remaining peptides in the series. There is, however no apparent correlation between ROS production and the reduction potentials as e.g. the peptide that is best a producing ROS (GHGGHG) has a similar reduction potential to the second worst (GHHG) (Fig. 3B, 5C).

### Binding geometry assessed by absorption spectroscopy

To investigate whether the differences in catalytic properties could be attributed to different chemical surrounding of the Cu(II) ion, we performed spectroscopic titrations using copper chloride for the series of peptides. We monitored the position of the weak d-d band in the copper complexes, as the absorption maxima (λ_max_) depends on the coordination mode in the peptide:Cu^2+^ complex. This parameter can be used to distinguish between 4N, 3N and 2N complexes: Truncated Aβ_4-16_ and GGH peptides coordinate Cu(II) in a 4N configuration with maximum absorption at 525 nm^14,34^. In contrast, 3N Cu(II) coordination results in a λ_max_ of around 600 nm as previously reported for GHK^35^, XHX^36^, and GHTD peptides^37^. Spectra with a λ_max_ near 660 nm are indicative for 2N complex GGH peptide^38^.

The peptide with a single histidine residue (GHG) had a λ_max_ of around 600 nm characteristic of 3N binding modes. Peptides with more than single His residue, separated by a single Gly residue, showed blue shift of maximum absorption to ~585 nm. Similar spectra were previously observed for N-terminally truncated Aβ_4-16_ and interpreted as Cu(II) binding between residues 11-16 in a 3N binding mode with a different coordination (amide nitrogen, two His residues)^14^. Without any spacing between His residues, or when the spacing was larger than one Gly, the maximum absorption showed red shift, suggesting different Cu(II) binding mode characterized by an increased content of 2N coordination when there are at least two intervening glycine residues. This suggests a binding mode where either the two nitrogen atoms from histidine residues coordinate the Cu(II) ion or a combination of a backbone amide and an imidazole side chain.

## Discussion

We have investigated the minimal requirements ROS production from Cu(II) binding disordered peptides using synthetic peptides. We found that ROS production occurs efficiently in peptides with two coordinating histidine side chains. An exact spacing is not required, suggesting that the intervening peptide chain remain flexible, but spectroscopic studies suggest a preference for complexes with a 2N geometry.

To be able to catalyze redox reactions, the peptide has to facilitate an “in-between state” where the copper ion can cycle between Cu(I)/(II). Our peptide suggest that this can be achieved simply by two histidine residues or one histidine and one cysteine close in the primary sequence. The lack of spacer length dependence indicates that the intervening sequence remains flexible, and the copper ion is mainly coordinated by the two histidine residues. This would imply that copper binding is common in disordered proteins. We do not explicitly test whether other residues are tolerated in the spacer between the histidine residues. However, as only glycine residues are used, this provides a neutral backbone that cannot form any side chain driven interactions with the copper-ion. This is likely true for several residue types in the intervening sequence, but not necessarily all. The preference for 2N geometries in ROS productions suggest that residues that provide additional coordination interactions with the copper ion may become inhibitory. Additionally, glycine is an unusual amino acid in having a highly flexible backbone. Other intervening residues may have conformational preferences that are inhibitory to forming a bi-dentate copper binding site. Nevertheless, it is likely that many other sequences with two closely spaced histidine residues and without inhibitory effects can form redox competent copper-binding sites.

Secondly, we set out to explore how the spacing between residues affect copper binding and ROS production. How “closely spaced” do histidine residues need to be? In multivalent interactions between IDPs the effective concentration of the second binding interaction follow a polymer scaling law with increasing linker length,^39,40^ and avidity enhancement follows.^41–43^ In bivalent protein complexes, avidity enhancement occurs when the effective concentration reaches the affinity of the monovalent interaction^43^. This predicts that the K_D_ approach a plateau close to the value recorded for a single histidine residue when the linker is very long - and by extrapolation the ROS production would decrease similarly. We do not seem to be close to this limit. However, as discussed above effective concentration might not be the right conceptual frame to describe the effects of multivalency when the individual interactor is a single residue. As a linker gets longer, the probability of introducing another side chain that coordinates weakly to the copper ion increases. The longest spacing considered here is seven residues, which correspond to the spacing in the octarepeat copper binding domain in PrP and the complex formed involving H6, H13 and H14 in Aβ. We see little reduction in ROS production up to such spacings are not inherently problematic for forming a copper:peptide complex competent for redox cycling – thus confirming using a minimal peptide series what was found previously in pathological proteins.

In conclusion, we find that the sequence motif for binding copper ions and catalyzing ROS production can be boiled down to two proximal histidine residues with a flexible spacing. This suggests that copper-binding and ROS production is not restricted to a small number of pathological proteins but may be a common property of intrinsically disordered proteins.

## Acknowledgements

This work was supported by the Novo Nordisk Foundation CO_2_ Research Center (CORC) with grant number NNF21SA0072700 to N.L. and M.K. and an instrument grant for the Carlsberg Foundation to Esben Lorentzen (CF22-0971).

