## Supplementary information for "Reactive oxygen generation by minimal copper binding peptide motifs"

#### Contents:

**Supplementary figures:**

Fig. S1-S4: Multi component fits of $G(HG)_x$ peptide series.

Fig. S5: ITC data for GCGGHG peptide

**Supplementary tables:**

Table S1-S4: Fitting parameters for multi-component fits of $G(HG)_x$ peptide series.

#### GHG peptide

**Figure S1:** Fitted thermograms for ITC experiments measuring binding of the GHG peptide to $CuCl_2$.Each panel represents a separate experiment.

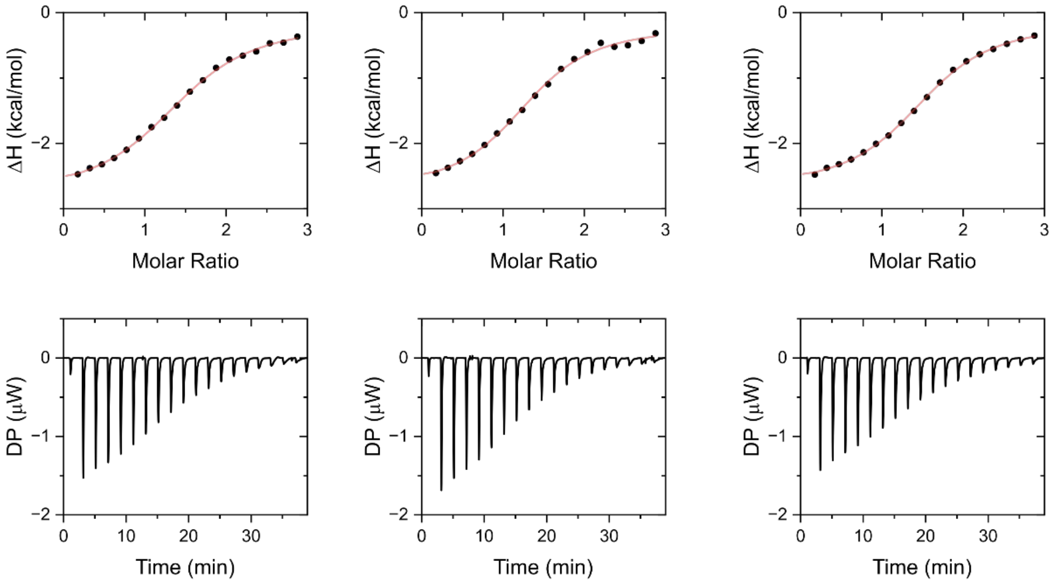


**Table S1:** Fitting parameters for ITC experiments measuring binding of the GHG peptide to $CuCl_2$. Each line corresponds to a fit to an independent experiment.

| Model | N | $K_D$ (μM) | ΔH (kcal/mol) |
|---|---|---|---|
| Single site | 1,48 | 37,0 | -2,64 |
| | 1,34 | 29,6 | -2,52 |
| | 1,55 | 28,4 | -2,54 |

### GHGHG peptide

**Figure S2:** Fitted thermograms for ITC experiments measuring binding of the GHGHG peptide to $CuCl_2$. Each panel represents a separate experiment.

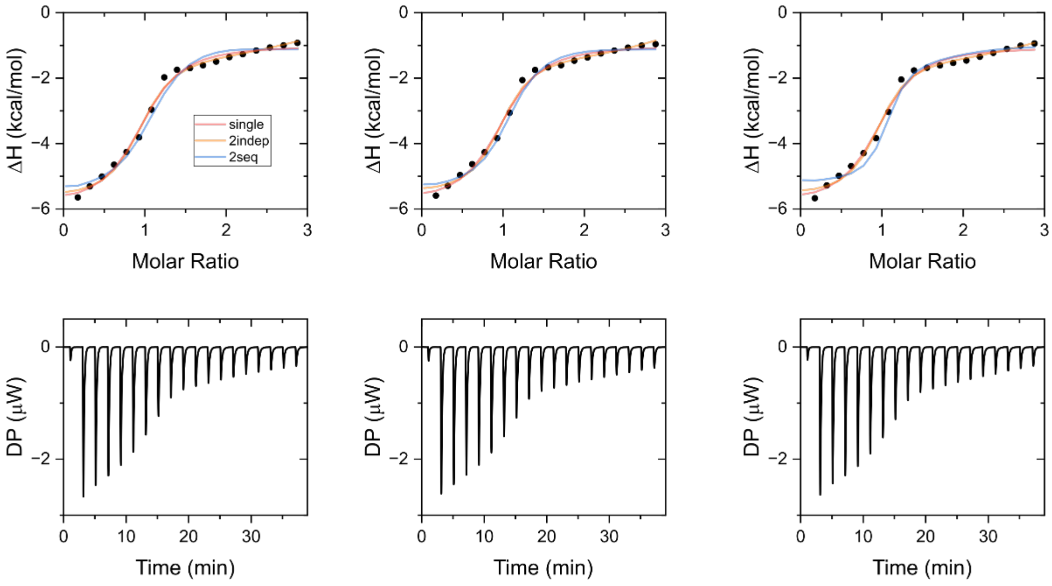


**Table S2:** Fitting parameters for ITC experiments measuring binding of the GHGHG peptide to $CuCl_2$. Each line corresponds to a fit to an independent experiment.

| **Model** | **N** | **$K_D$ (μM)** | **ΔH (kcal/mol)** |
|---|---|---|---|
| Single site | 0,965 | 13,2 | -4,82 |
| | 0,959 | 13,8 | -4,91 |
| | 0,978 | 13,6 | -4,83 |

| | **$N_1$** | **$K_{D1}$ (nM)** | **$\Delta H_1$ (kcal/mol)** | **$N_2$** | **$K_{D2}$ (μM)** | **$\Delta H_2$ (kcal/mol)** |
|---|---|---|---|---|---|---|
| 2 independent sites | 0,905 | 397 | -5,54 | 2,34 | 21,8 | -1,3 |
| | 0,898 | 537 | -5,64 | 2,37 | 24,9 | -1,22 |
| | 0,919 | 401 | -5,64 | 2,36 | 22,1 | -1,44 |

| | **$K_{D1}$ (nM)** | **$\Delta H_1$ (kcal/mol)** | **$K_{D2}$ (μM)** | **$\Delta H_2$ (kcal/mol)** |
|---|---|---|---|---|
| Sequential binding of 2 | 414 | -4,15 | 25,3 | -0,544 |
| | 386 | -4,22 | 7,61 | 0,058 |
| | 475 | -4,16 | 12,1 | -0,029 |

#### GHGHGHG peptide

**Figure S3:** Fitted thermograms for ITC experiments measuring binding of the GHGHG peptide to $CuCl_2$. Each panel represents a separate experiment.

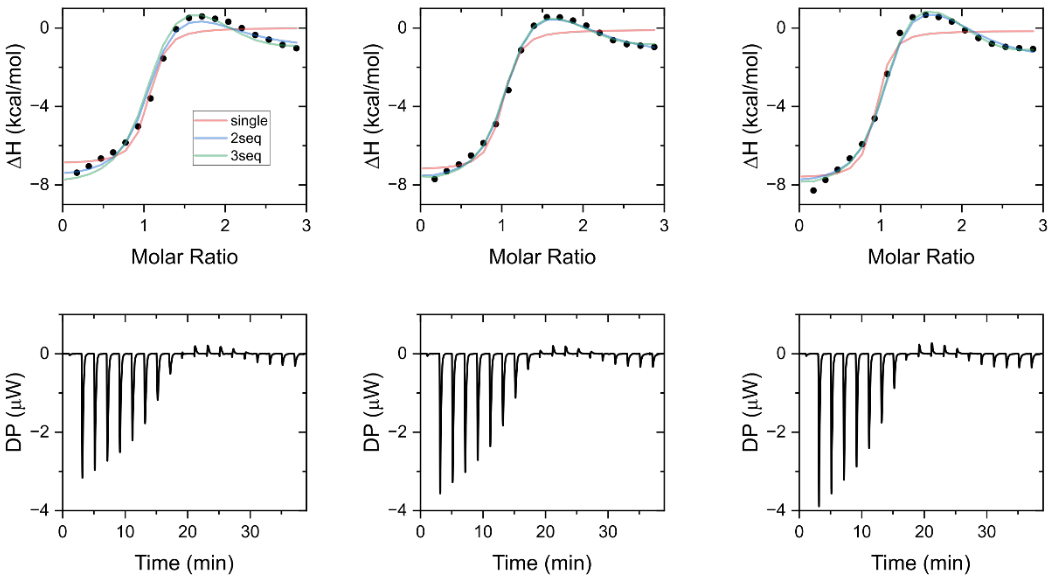


**Table S3:** Fitting parameters for ITC experiments measuring binding of the GHGHG peptide to $CuCl_2$. Each line corresponds to a fit to an independent experiment.

| **Fitting model** | **N** | **$K_D$ (µM)** | **ΔH (kcal/mol)** | | | |
|---|---|---|---|---|---|---|
| Single site | 0,89 | 1,86 | -7,51 | | | |
| | 1,00 | 2,06 | -6,93 | | | |
| | 0,955 | 2,18 | -7,16 | | | |
| | **$K_{D1}$ (nM)** | **$\Delta H_1$ (kcal/mol)** | **$K_{D2}$ (µM)** | **$\Delta H_2$ (kcal/mol)** | | |
| 2 sequential sites | 766 | -6,00 | 29,7 | 4,63 | | |
| | 830 | -6,32 | 31,9 | 3,04 | | |
| | 815 | -6,14 | 34,0 | 3,82 | | |
| | **$K_{D1}$ (nM)** | **$\Delta H_1$ (kcal/mol)** | **$K_{D2}$ (µM)** | **$\Delta H_2$ (kcal/mol)** | **$K_{D3}$ (µM)** | **$\Delta H_3$ (kcal/mol)** |
| 3 sequential sites | 224 | -7,89 | 7,02 | 2,76 | 80,0 | -3,95 |
| | 300 | -7,74 | 8,70 | 2,5 | 98,1 | -3,57 |
| | 267 | -7,67 | 9,11 | 2,2 | 76,9 | -3,13 |

#### GHGHGHGHG peptide

**Figure S4:** Fitted thermograms for ITC experiments measuring binding of the GHGHG peptide to $CuCl_2$. Each panel represents a separate experiment.

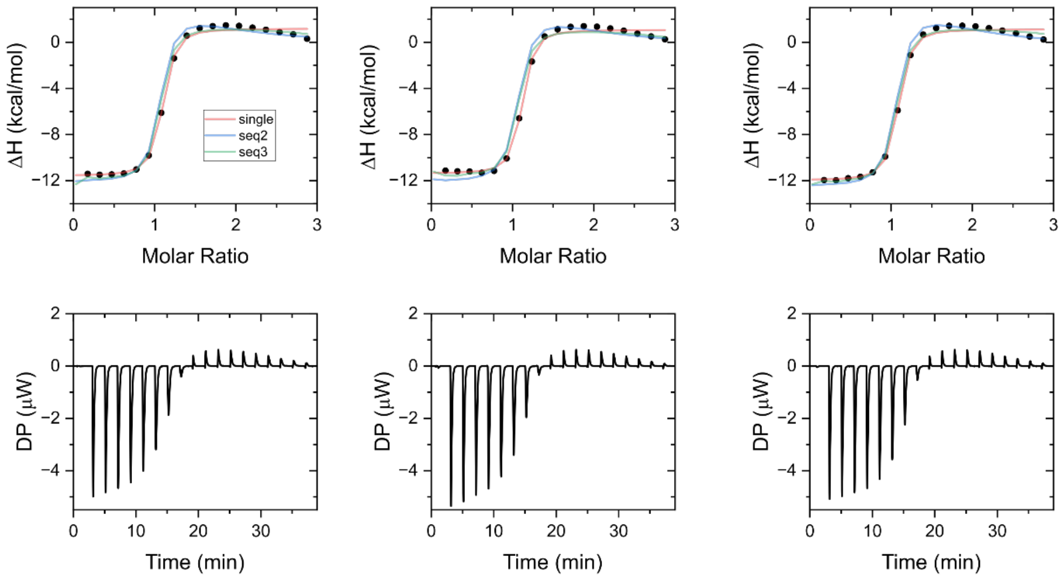


**Table S4:** Fitting parameters for ITC experiments measuring binding of the GHGHG peptide to $CuCl_2$. Each line corresponds to a fit to an independent experiment.

| | **N** | **$K_D$ (nM)** | **ΔH (kcal/mol)** |
|---|---|---|---|
| Single site | 1,04 | 1010 | -12,8 |
| | 1,05 | 846 | -12,4 |
| | 1,02 | 993 | -13,1 |

| | **$K_{D1}$ (nM)** | **$ΔH_1$ (kcal/mol)** | **$K_{D2}$ (μM)** | **$ΔH_2$ (kcal/mol)** |
|---|---|---|---|---|
| Sequential binding of 2 | 345 | -12,0 | 67,7 | 3,15 |
| | 338 | -11,9 | 58,8 | 3,07 |
| | 369 | -12,2 | 67,0 | 3,79 |

| | **$K_{D1}$ (nM)** | **$ΔH_1$ (kcal/mol)** | **$K_{D2}$ (μM)** | **$ΔH_2$ (kcal/mol)** | **$K_{D3}$ (μM)** | **$ΔH_3$ (kcal/mol)** |
|---|---|---|---|---|---|---|
| 3 sequential sites | 54,0 | -11,8 | 9,42 | 1,37 | 35,1 | 1,44 |
| | 51,9 | -11,2 | 9,39 | 1,50 | 33,9 | 1,50 |
| | 55,2 | -12,1 | 9,41 | 1,35 | 25,0 | 1,24 |

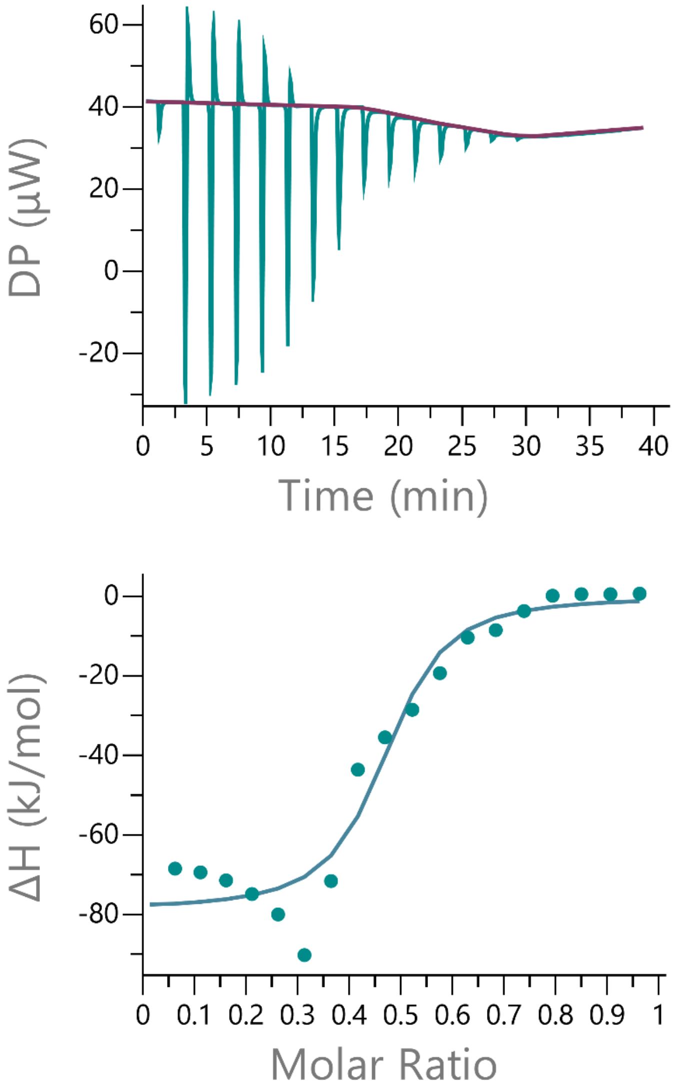


**Figure S5: ITC** experiment titrating 3mM $CuCl_2$ into 600 µM GCGGHC peptide. The experiment does not resemble any standard binding model and was not interpreted further.
